# mm2-gb: GPU Accelerated Minimap2 for Long Read DNA Mapping

**DOI:** 10.1101/2024.03.23.586366

**Authors:** Juechu Dong, Xueshen Liu, Harisankar Sadasivan, Sriranjani Sitaraman, Satish Narayanasamy

**Author notes:** Both authors contributed equally to this research (Juechu Dong, Xueshen Liu).

## Abstract

Long-read DNA sequencing is becoming increasingly popular for genetic diagnostics. Minimap2 is the state-of-the-art long-read aligner. However, Minimap2’s chaining step is slow on the CPU and takes 40-68% of the time especially for long DNA reads. Prior works in accelerating Minimap2 either lose mapping accuracy, are closed source (and not updated) or deliver inconsistent speedups for longer reads. We introduce *mm2-gb* which accelerates the chaining step of Minimap2 on GPU without compromising mapping accuracy. In addition to intra- and inter-read parallelism exploited by prior works, *mm2-gb* exploits finer levels of parallelism by breaking down high latency large workloads into smaller independent segments that can be run in parallel and leverages several strategies for better workload balancing including split-kernels and prioritized scheduling of segments based on sorted size. We show that *mm2-gb* on an AMD Instinct™ MI210 GPU achieves 2.57-5.33x performance improvement on long nanopore reads (10kb-100kb), and up to 1.87x performance gain on super long reads (100kb-300kb) compared to SIMD accelerated mm2-fast. *mm2-gb* is open-sourced and available at https://github.com/Minimap2onGPU/mm2-gb.

## 1 Introduction

Long-read DNA sequencing offers the unique advantage of being able to span highly repetitive regions in the genome, thereby, enabling applications such as structural variant calling as well as de novo assembly [7, 15]. A recent study demonstrated the world’s fastest blood-to-variants workflow using long-read sequencing to perform genetic diagnosis at the point of care [7]. Oxford-Nanopore Technology (ONT) [4] and Pacific Biosciences (PacBio) [2] are the two major platforms for long-read sequencing, and their accuracy, throughput and read lengths are improving with rapid advancements in single-molecule sequencing technology. Jain et al. [9] reported ultra-long reads up to 882kb (N50 > 100kb) that enable assembly and phasing of 4-Mb Major Histocompatibility Complex (MHC) locus in its entirety.

Minimap2 [11] is the state-of-the-art long read mapper and aligner that maps DNA or long mRNA sequences against a reference database. Even though Minimap2 is well-engineered to handle noisy long reads 100kb, its CPU implementation struggles to keep up with the growing sequencing throughput (>1Tbp per day [18]). In particular, the chaining kernel in the seed-chain-align strategy of Minimap2 is an increasingly pronounced bottleneck for ultra-long reads. Time spent in chaining grows from 40% to 68% as average ONT read length grows from 2.25Kb to 127.11Kb [16]. This motivates us to accelerate Minimap2 for higher throughput on a GPU.

Unlike accelerating base-level alignment on GPU [6, 19], the chaining kernel is highly irregular and difficult to accelerate in terms of finding enough parallelism and balancing the workload. Prior attempts to accelerate chaining on GPU either sacrifice mapping accuracy [8] or suffer from diminishing benefits in long reads >20kb (mm2-ax [16]).

In this study, we introduce Minimap2-gigabases (*mm2-gb*), an enhancement over the mm2-ax through the incorporation of three optimization strategies aimed at augmenting intra-read parallelism and achieving more equitable workload distribution on GPUs. These optimizations include: segment cutting (Sec. 3.1), split score generation kernels (Sec. 3.2), and prioritized long segment scheduling (Sec.3.3.2). We break reads into independent segments to leverage intra-read parallelism. We configure three GPU kernels to process segments of different lengths separately, thus improving workload balancing within each kernel. We implement prioritized long-segment scheduling to improve inter-block workload balance. To this end, *mm2-gb* delivers a consistent speedup of 2.57-5.33x for chaining of coarse-grained long ONT reads (10kb-100kb) on AMD Instinct™ MI210 GPU [1] compared to mm2-fast, a vectorized version of Minimap2 running on 32 Intel® 1 Icelake cores with AVX-512 extension. *mm2-gb* delivers up to 1.87x speedup on ultra-long ONT reads (100kb-300kb). Further, in Sec. 8 we discuss future work to potentially improve performance for ultra-long reads. *mm2-gb* supports HIP and CUDA runtimes and is open-sourced.

## 2 Background

### 2.1 Overview of Minimap2

Minimap2 is the most popular DNA/mRNA long sequence mapper and aligner for popular long-read sequencing platforms like Oxford Nanopore Technologies (ONT). It aims to process ultra-long reads of 100Kb on average at high throughput. Minimap2 performs long sequence mapping in three steps: seeding, chaining, and alignment (Fig. 1). Seeding identifies exact matches of short k-base minimizers (**anchors**) between the query and reference sequence. Seeding is fast because the references are pre-processed offline and indexed to a multimap hash table for quick lookup. Chaining takes anchors sorted by reference position as input and identifies sub-regions of the target reference that roughly map to each other. Minimap2 solves chaining as a 1-D dynamic programming problem. For workloads such as full genome or assembly alignment that requires base-level alignment, Minimap2 runs an alignment kernel using Needleman-Wunsch [13] with Suzuki-Kazahara [17] formulation to close the gaps between adjacent anchors in the chains. These gaps are usually short and are solved as a 2D dynamic programming problem.

**Figure 1.**
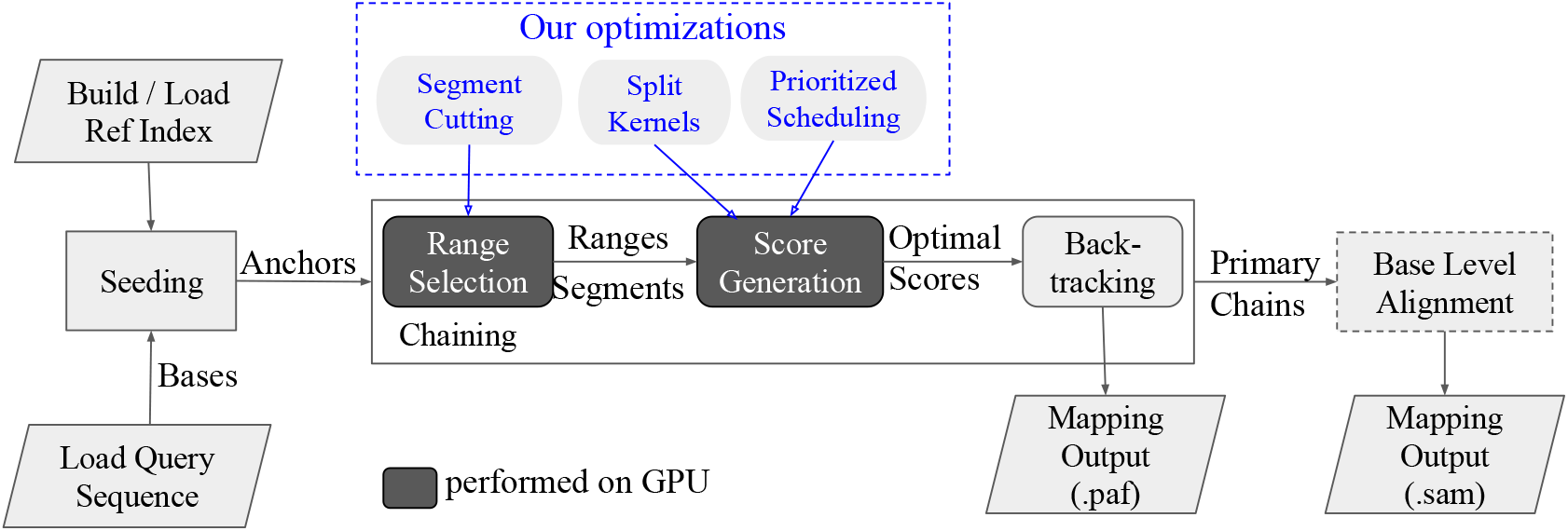
Minimap2 Workflow: Steps of chaining performed on the GPU is shown in dark grey blocks. Our optimizations to GPU kernels are shown in the blue box.

### 2.2 Chaining is the Bottleneck

The chaining process takes sorted anchors as input and performs 1D dynamic programming to identify subsets of ordered anchors that are in linear alignment on the reference and query sequence. This is achieved by 1D-dynamic programming with a simple cost function: by sequentially going through the read, each anchor attempts to chain against its predecessor anchors that are less than *max_dist_x* bases apart on the reference sequence and selects the predecessor that generates the optimal chaining score. The chaining score is calculated as the optimal score of the predecessor plus a reward based on the gap between the anchor pair on the query and reference sequence. After generating optimal scores for all the anchors, Minimap2 performs backtracking and identifies primary chains. Among all the steps of chaining, the optimal score calculation is the most compute-intensive (>95% of chaining CPU time).

For ease of understanding, we define the following terminologies:

- *Distance between 2 anchors* (in bases): The number of bases between two anchor locations on the reference read. Each anchor attempts to chain against all anchors within the *max_dist_x* distance.
- *Predecessor/Successor Range* (in # of anchors): The number of preceding/succeeding anchors within*max_dist_x*of a given anchor.

### 2.3 Quadratic Complexity of Chaining

Prior works [8, 10, 16] have found chaining to be the bottleneck in aligning long ONT reads with Minimap2 – 40% to 70% of CPU time is spent in chaining. The chaining kernel’s complexity is quadratic with respect to read length while the seeding algorithm scales linearly. Additionally, recent advances in sequencing technologies promise ultra-long reads of 100kb on average with high throughput [9]. Therefore, it is fair to argue that chaining would become an even bigger bottleneck.

To perform a fair evaluation of chaining, we evaluate the performance of chaining kernels in chaining scores calculated per second, instead of anchors processed per second.

The compute-intensive step of chaining calculates the score between every anchor pair that are within a predefined distance on the reference sequence. Because the number of anchors within this distance is varied, anchors are not distributed evenly on the reference sequence – they are denser in regions where the reference maps well to the query. With higher anchor density, an anchor in these regions finds more predecessors within *max_dist_x* bases, leading to significantly more anchor pairs to compute scores with during chaining. To demonstrate the effects of uneven distribution, Fig. 2 shows that the average amount of score generations grows with read length, because there are more compute-dense regions in long reads, while the anchors per base remains stable. We argue that using scores generated per second instead of anchors per second as the metric of chaining evaluation reflects real performance more accurately.

**Figure 2.**
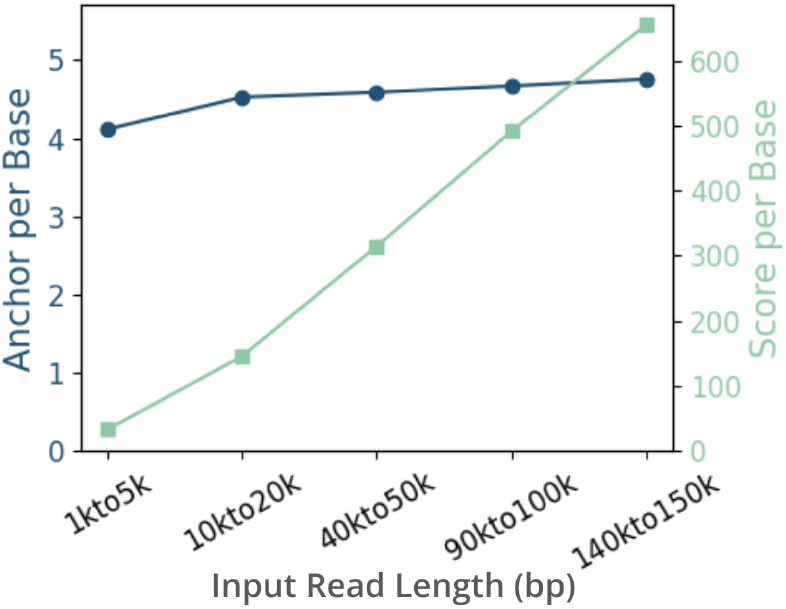
Longer reads have more chain scores to be generated per base while the total number of anchors per base remains almost the same. X-axis shows query datasets of different read lengths.

## 3 Design

In this work, we design *mm2-gb*, a GPU chaining kernel for Minimap2 that does not compromise on mapping accuracy and delivers consistent performance on reads of various lengths, including ultra-long reads (*>*100kb). We show that *mm2-gb* delivers consistent speedup on coarse-grained long reads (10kb-100kb) compared to mm2-fast [10], a SIMD-vectorized version of Minimap2 on 32 Intel®^1^ Icelake cores with AVX-512 extension. Another prior work, mm2-ax [16], relies on CPU to pre-process the reads and provides decent speedups on GPU for reads <20kb, but fails to deliver consistent speedups for longer reads, where chaining becomes a more pronounced bottleneck due to its quadratic compute complexity. We introduce three major optimizations to improve intra-read parallelism and workload balancing: segment cutting, split score generation kernels, and prioritized long segment balancing as shown in Fig. 3. Segment cutting breaks the sequence of sorted anchors into independent segments and allows *mm2-gb* to process them in parallel. We split the computationally intensive score generations kernel into three sub-kernels, each specialized in handling short (<2k anchors), mid (2k-10k anchors), or long (>10k anchors) segments. We then enhance the long segment score generation kernel with prioritized ultra-long segment scheduling to improve workload balance between GPU thread blocks.

**Figure 3.**
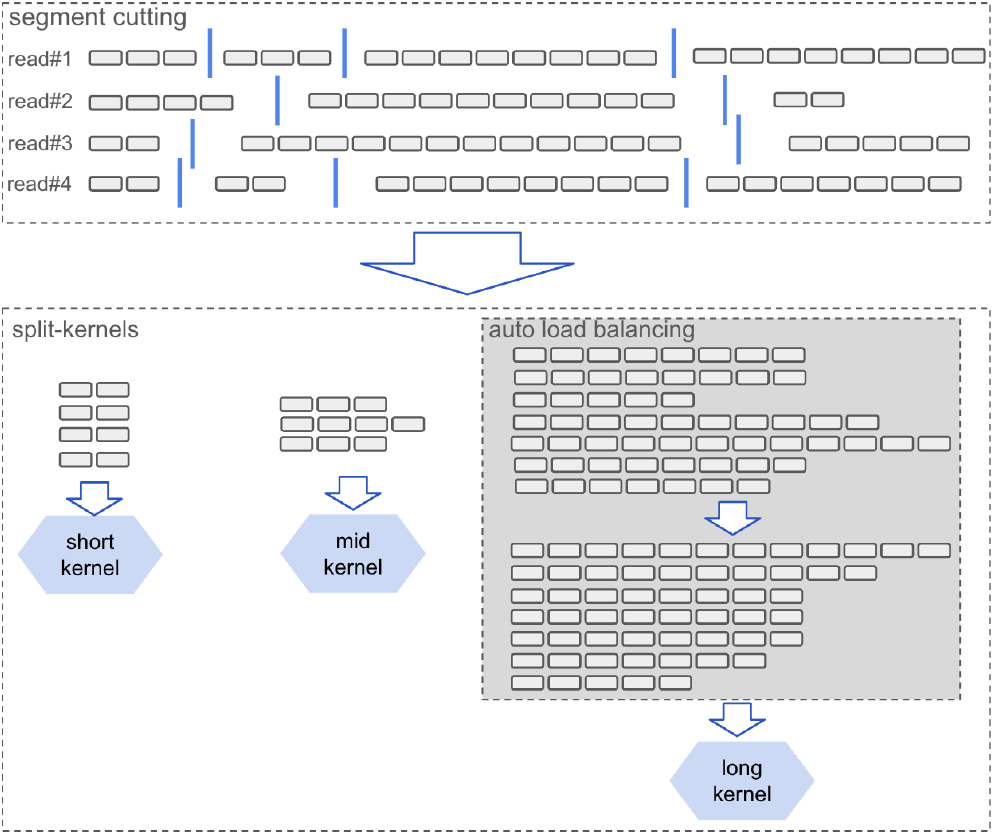
*mm2-gb* optimizations: segment cutting, split-kernels and auto-load-balancing

To this end, we build a range selection kernel on GPU that performs forward range selection and identifies independent segments within a read. Independent segment identification eliminates the need to sequentially process all the anchors in a read with a single thread block on the GPU and instead enables multiple blocks to work on different segments of a read in parallel. The split-kernel design helps us navigate the difficulty in launching a single kernel that handles workloads of different sizes in a balanced manner. Instead, the split-kernel method relies on three independent kernels optimized to better handle segments of different compute densities (short, medium, and long). Finally, we improve workload balancing by applying atomic operation-based prioritized scheduling and long segment aggregation to amortize the latency of ultra-long segments.

We compare our GPU-accelerated Minimap2 (*mm2-gb*) on GPU to SIMD-accelerated mm2-fast by mapping real ONT reads to human genome reference. We show that *mm2-gb* generates identical mapping results as mm2-fast and delivers 2.57x-5.33x speedup on coarse-grained long reads (10kb - 100kb). *mm2-gb* supports HIP and CUDA environments on AMD and NVIDIA GPUs. *mm2-gb* is an open-sourced and accelerated implementation of Minimap2 without compromising mapping accuracy.

### 3.1 Improve GPU Occupancy by Breaking Down Reads into Segments

To maximize the occupancy on data center GPUs such AMD Instinct™ MI210 which can run 208k threads in parallel, it is crucial to identify sufficient parallelism in the optimal score generation step of chaining. Similar to mm2-ax, *mm2-gb* adapts its forward loop transformation technique and leverages inter-read and intra-range parallelism at the grid and block levels, respectively, in the score generation kernel. Intra-range parallelism arises from threads within a block performing forward score generation between the successors within range against a given anchor. Inter-read parallelism emerges from the data parallelism between different reads.

To address the lack of parallelism in long ONT reads (>20kb) and achieve higher GPU occupancy, *mm2-gb* introduces a new level of parallelism, intra-read parallelism, to long-read chaining. Instead of serializing the optimal score generation for the sorted anchors belonging to the same read, *mm2-gb* divides the anchors into multiple independent segments and processes them in parallel. This is done carefully to avoid violating data dependencies between finding the optimal chain score of an anchor and its possible predecessors. mm2-ax [16] demonstrated that 67% of the input anchors have a successor range of zero and do not start a chain. We observe that the anchors succeeding such a zero-range anchor are too distant to chain against it or any of its predecessors. Therefore, the chain scores for anchors succeeding the zero-range anchor are not dependent on its predecessor scores or itself. The sorted input anchors are cut at these zero-range positions and divided into segments of anchors that are then processed independently. We perform segment cutting in the successor range selection kernel on GPU (Algorithm 1).

Our parallel segment cutting algorithm is designed to divide reads into short segments, each approximately the size of one range selection block (512 in our evaluated configuration). In the range selection kernel, each thread is tasked with calculating the successor range for one anchor in the read. The last few threads in each block (10 threads in our evaluated configuration) attempt to update the cut position with their anchor index if they encounter a zero-range anchor. Consequently, our algorithm effectively partitions reads into segments of 512 anchors, unless none of the last 10 threads encounters a zero-range anchor. This algorithm introduces minimal overhead to the range selection kernel and yields segments with an average length of 546.9 anchors on a subset of the 60x HG002 ONT dataset (average read length 45kb).

### 3.2 Split Score Generation Kernels: Specialize Kernels for Different Range Profiles

In mm2-ax [16], Sadasivan et al. report that the chaining kernel has highly irregular arithmetic intensity and intra-range parallelism, characterized by a high variance in the successor range. We extend this idea by measuring workload in terms of chaining scores computed to reflect the mapping cost more accurately. The statistics on a dataset subset from 60x HG002 ONT [5] shows that 93.5% of anchors have a range less than 64, but they only form less than 2% of the anchors-pairs that require a score calculation. The successor ranges of the remaining 6.5% anchors follow a long-tail distribution (Fig. 4a). The score calculation workload is distributed across anchors of different range profiles (Fig. 4b), and exhibits different arithmetic intensity and intra-range parallelism. It is hard to design one kernel that suits all the anchors with different range profiles.

**Figure 4.**
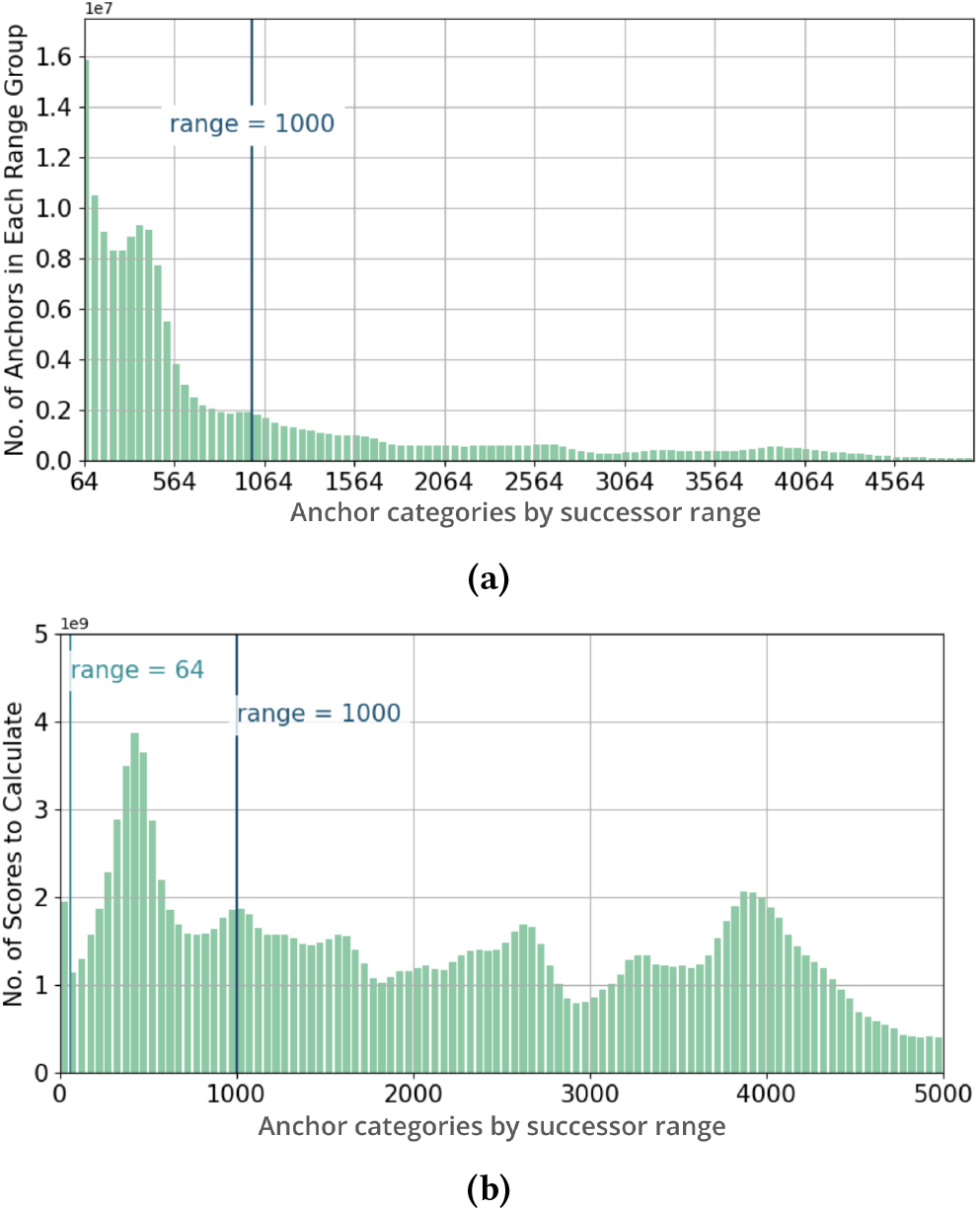
Successor range distribution is highly imbalanced. (A)Anchors with a range >64 follow a long-tail distribution (B)Workload is broadly distributed among anchors of different range profiles.

#### Algorithm 1

Segment Cutting In Range Selection Kernel

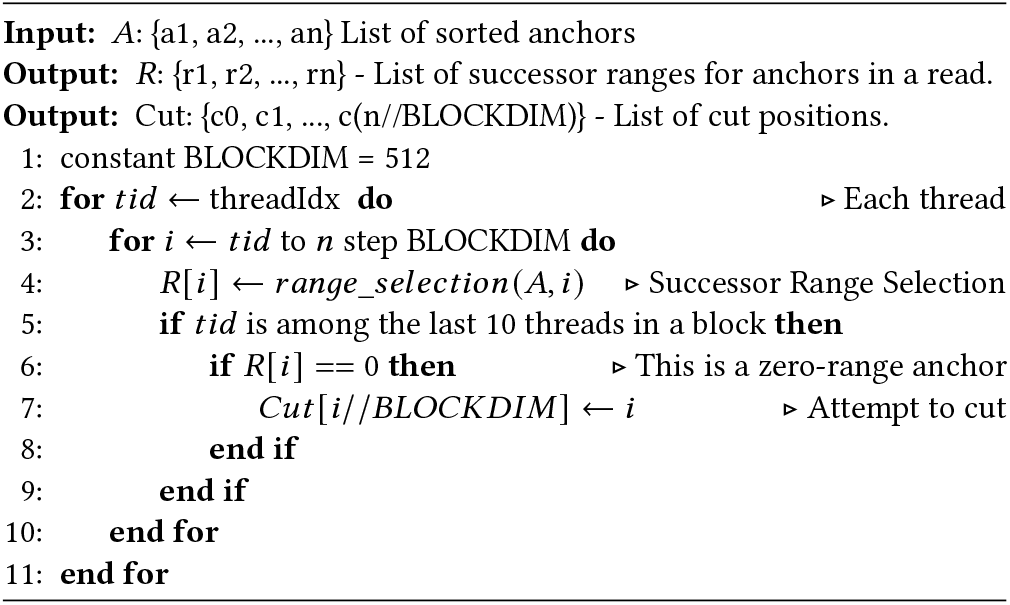

To overcome this irregularity, we develop three separate kernels, short, mid, and long, to handle anchors of various ranges. We distribute anchors into the three different kernels based on the correlation between the segment length and average anchor range.

The short kernel is optimized for anchors with a range shorter than the wavefront size (<64 threads). Each block in the short kernel contains only one wavefront that is executed in lock-step to avoid block synchronization between iterations. This also allows us to fit a larger grid dimension on the device to better leverage the inter-segment parallelism. The mid-kernel is optimized for anchors with more intra-range parallelism to benefit from a larger block size of 512. The long kernel is optimized for anchors with long ranges (>1024) and needs 1024 threads to work on forward score generation in parallel.

Through our analysis of the range profile of segments, we find that anchors in longer segments tend to have a larger average range (Fig. 5a). Because segment is the minimum parallel unit due to succeeding dependency, by assigning segments into short, mid and long kernels based on their length, anchors are distributed into appropriate chaining kernels. From a zoomed-in view (Fig. 5b), we find segments shorter than 2k anchors have an average range of less than 64 and are optimal workloads for the short kernel. *mm2-gb* assigns the segments of medium length (2k-10k anchors) to the mid kernel and leaves segments in the long tail to the long kernel (Fig. 6). Note that the long kernel is responsible for generating 86% of the scores, and therefore an optimized long kernel is the key to achieving high chaining throughput.

**Figure 5.**
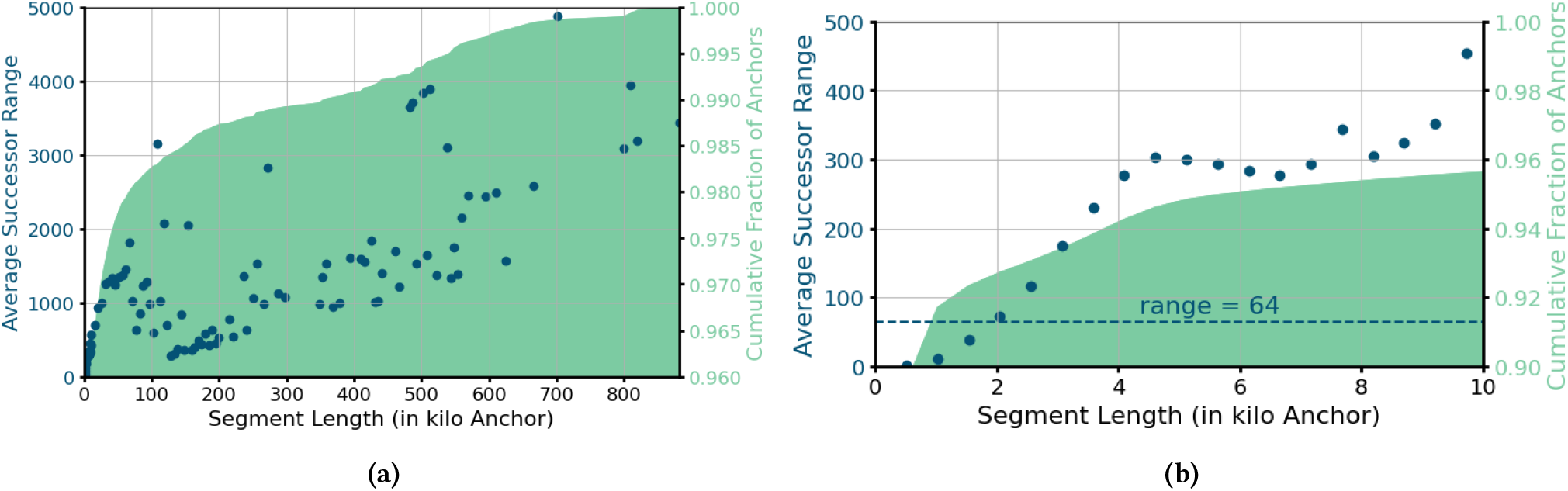
Long segments consist of anchors with larger range profiles. Segment Length is the number of anchors in each segment. (a) The number of anchors in long segments follows a long tail distribution up to 1 million. (b) Zoom-in View (segments <20k anchors): Segments less than 2k anchors long have an average range less than 64.

**Figure 6.**
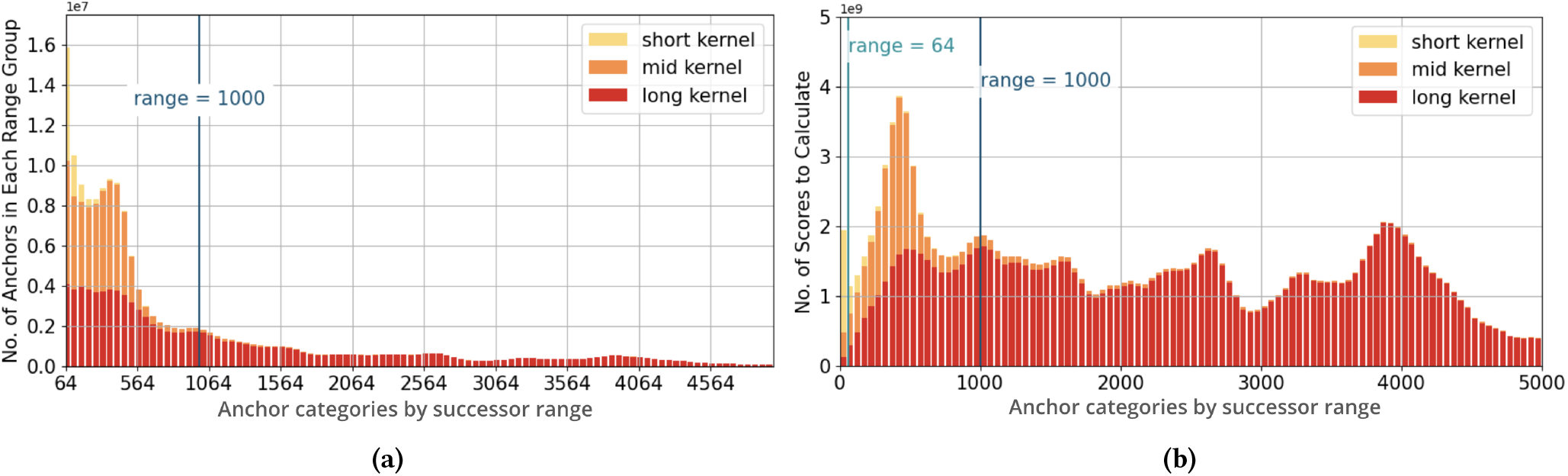
Anchors of different range profiles are assigned to their respective kernels. (a) Most anchors assigned to the mid kernel have a range <512. Half of the anchors between 64 to 512 in range are assigned to the long kernel. (b) The long kernel handles a majority (86%) of total work.

### 3.3 Long Kernel Scheduling

The runtime of the long kernel is bounded by the latency of processing ultra-long segments that can be up to 1 million anchors long. There are only a few ultra-long segments in a batch, but anchors in each ultra-long segment are handled sequentially by a single compute unit (CU). *mm2-gb* implements long segment aggregation and prioritized segment scheduling to improve the overall throughput of the long kernel in the presence of ultra-long segments.

#### 3.3.1 Long Segment Aggregation

The key to improving throughput for the long kernel is to identify enough long segments to fully occupy all 104 CUs on an AMD Instinct™ MI210 GPU. Reads are processed in batches sized to fit the 64 GB of High Bandwidth Memory (HBM) capacity in an MI210 GPU. From this batch, a small proportion of segments are long. The occupancy of the long kernel processing this small set of long segments from one batch is very low, only around 20.2%. To remedy this, we aggregate enough long segments from multiple input batches and launch a single long kernel to process all of them (Fig. 7 below). A buffer sized 15% of MI210’s HBM capacity can hold aggregated long segments from 4-5 batches of reads, and increase the CU occupancy to 85.5%.

**Figure 7.**
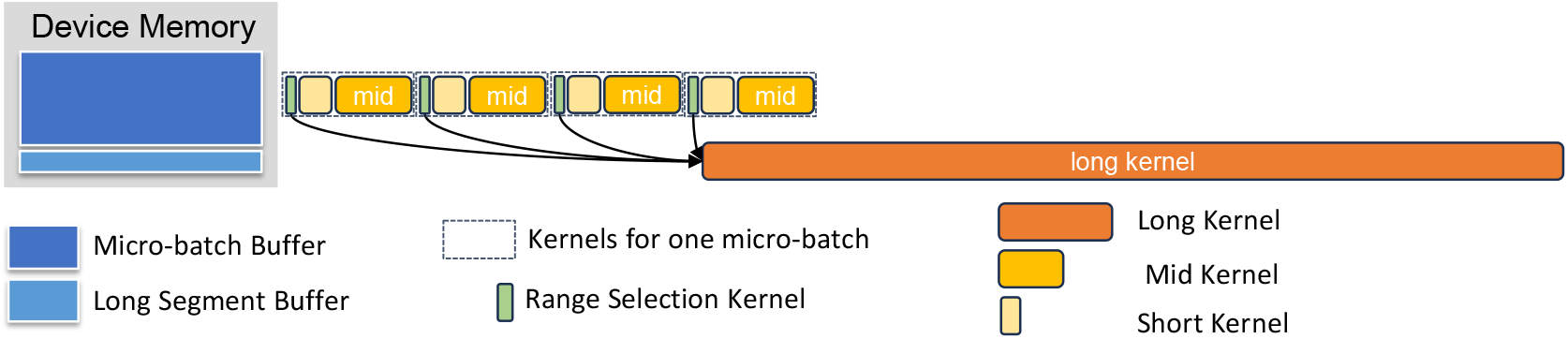
Aggregating long segments from multiple micro-batches into one long score generations kernel.

#### 3.3.2 Prioritized Segment Scheduling

In the long kernel, segments in descending order of lengths are scheduled to persistent blocks that remain active throughout the kernel runtime. This helps improve workload balance between blocks. After aggregating long segments from multiple batches of reads, *mm2-gb* sorts them by length in descending order. As proposed in Algorithm 2, once launched, each block in the long kernel pops the longest segment from the queue, processes it, and then pops the next in the queue to process until the queue is empty. This algorithm guarantees that: 1) Latency-bounded ultra-long segments are scheduled first. 2) Blocks that complete processing a segment early automatically get scheduled more work. As illustrated below in Fig. 8, prioritized segment scheduling significantly improves block-level workload balance in the long score generation kernel.

**Figure 8.**
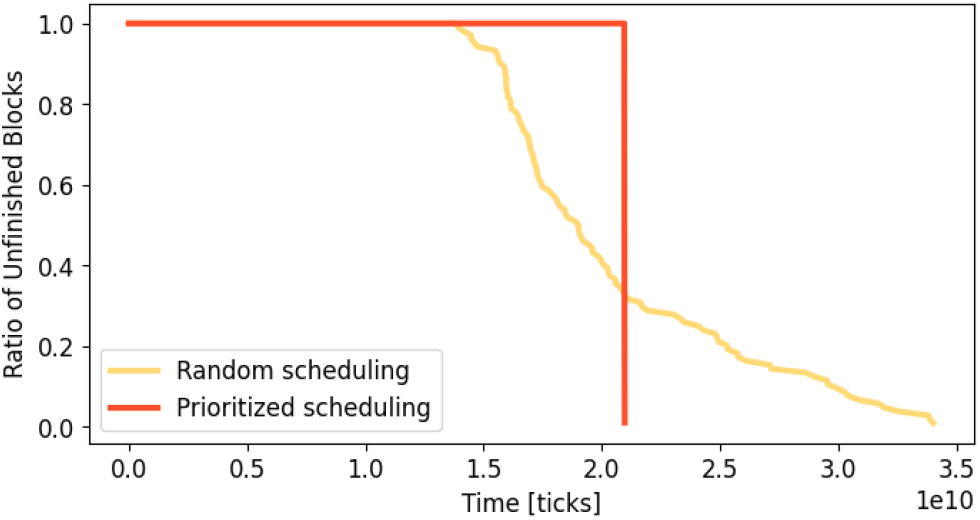
Inter-block workload balance of long score generation kernel with and without prioritized segment scheduling. Prioritized scheduling significantly reduces tail latency. The unit of time here is represented by tick count of per-multiprocessor counter.

##### Algorithm 2

Prioritized Segment Scheduling In Long Score Generation Kernel

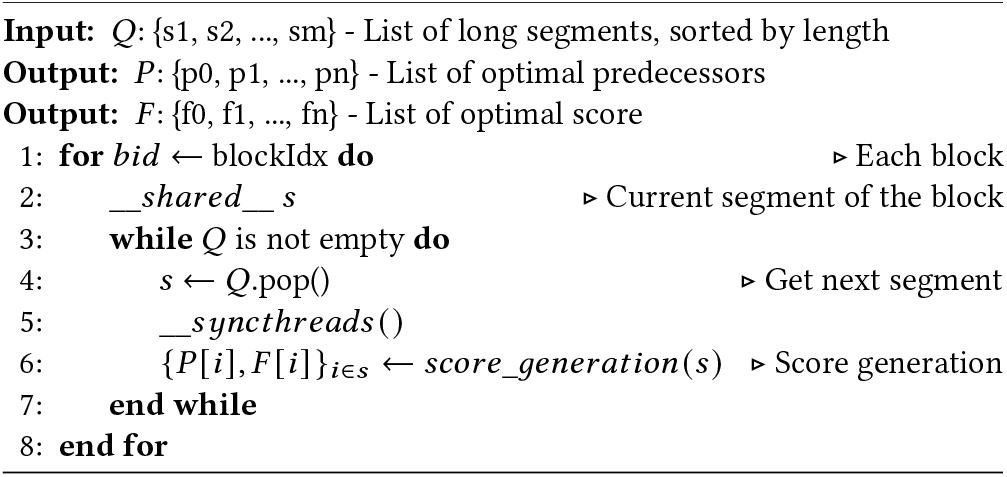

## 4 Evaluation

### 4.1 Experiment Setup

We accelerate Minimap2 v2.24, which is the same version used by mm2-fast [10]. The previous work, mm2-ax, is tied to an older version of mm2-v2.17, and is built to work on an NVIDIA A100 GPU. Given that v2.17 is less accurate and less computationally intensive compared to v2.24 [12], and mm2-ax is close-sourced to be tweaked, we didn’t use it as our baseline. Instead, we use SIMD accelerated mm2-fast and Minimap2 on GPU without our optimization (see Fig. 1) as baselines. Because mm2-ax also uses mm2-fast as its baseline, we are still able to compare *mm2-gb* with mm2-ax qualitatively. We demonstrate the benefits of our optimizations using AMD Instinct™ MI210 GPU on the AMD Accelerator Cloud (AAC) cluster [3]. Our performance is evaluated on a single AAC node containing 512GB of host memory and one AMD Instinct™ MI210 GPU with 64GB of HBM memory. We run mm2-fast on a server with 125GB memory and a 32-core Intel®^1^ Xeon®^2^ Gold 6326 CPU, which supports AVX512. Our test data is extracted from Oxford Nanopore Technologies (ONT)’s 60X HG002 dataset [5]. To demonstrate our performance gain on different read lengths, especially long reads, we allocate five bins based on read lengths (1k-10k, 10k-20k, 40k-50k, 90k-100k, and 100k-300k bases) and assign 5Gb of reads into each of them (similar to coarse-grained workload in mm2-ax [16]).

### 4.2 Summary of Results

Experiment results in Fig. 9 show *mm2-gb* achieves 2.57-5.33x speedup over mm2-fast in processing long reads (10k-100k) and up to 1.87x for super long reads (100kto300k). These coarse-grained long-read datasets are challenging to be parallelized on the GPU and mm2-ax achieves speedups downwards of 2x over mm2-fast, without the manual grouping of reads. We achieve 4.36x speedup on short reads (1kto10k), compared to up to 10x for mm2-ax. Potential reasons for the slowdown are discussed later in Sec. 7.

**Figure 9.**
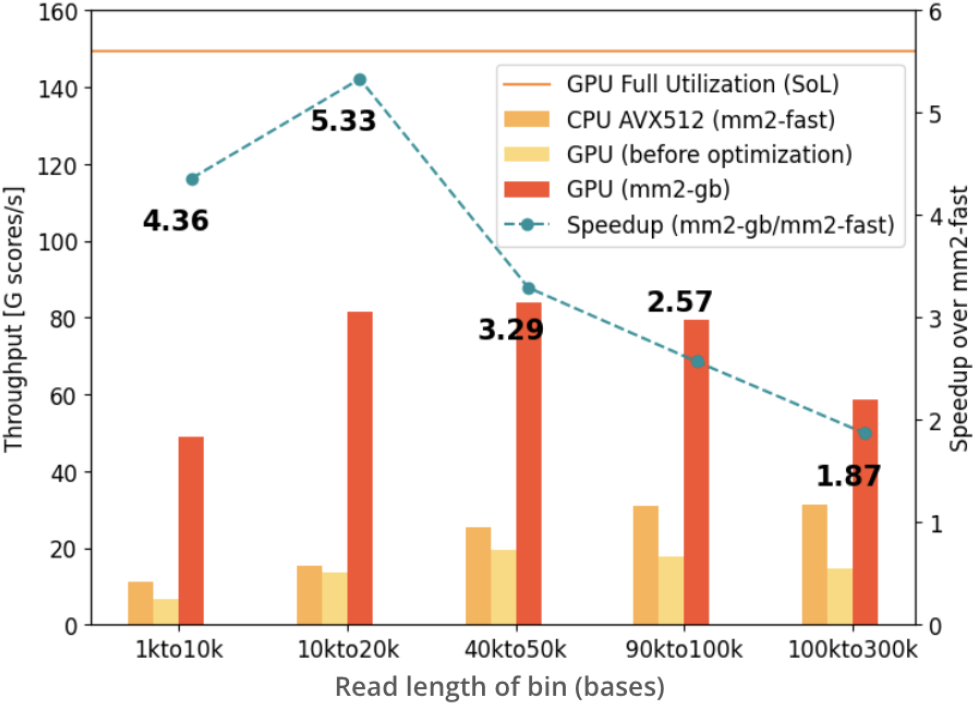
Maximum throughput in terms of scores generated per second, where *mm2-gb* achieves 1.87-4.36x speedup over mm2-fast.

Our *mm2-gb* on GPU maintains 50-60% of theoretical max throughput in processing long reads (10k-100k), and achieves 39% theoretical max throughput on super long reads (100k-300k), while before optimization it was at less than 15%. For short reads (1k-10k), because the low average range leads to a low workload intensity, we observe a 33% theoretical max throughput.

Accelerating chaining for mapping long DNA reads is a challenging task on GPU because of its long latency and low hardware occupancy. The results demonstrate that our efforts in splitting workload and improving workload balance makes a big difference. Compared with previous works like mm2-ax, our acceleration doesn’t require users to manually assign short reads to run concurrently with long reads (i.e. generate fine-grained datasets) to saturate the occupancy when mapping long read datasets, instead, by introducing the finer levels of parallelism in the segment, *mm2-gb* automatically schedules segments of different lengths to ensure the workload are distributed equally on each CU.

In summary, *mm2-gb* overcomes the latency issue in processing long and super-long reads, and achieves significant improvement over SIMD-accelerated mm2-fast and previous GPU-accelerated versions of mm2.

## 5 Conclusion

Chaining kernel performance is crucial for constructing a high-throughput long-read aligner capable of adapting to the increasing read lengths. However, it remains slow in the state-of-the-art long-read aligner Minimap2, consuming 40-68% of the total runtime. To address this bottleneck, we present Minimap2-gigabases (*mm2-gb*), a GPU-accelerated version of Minimap2 tailored for high throughput long-read chaining while maintaining the same accuracy. *mm2-gb* improves GPU workload balance and occupancy within the irregular chaining kernel by cutting long reads into independent segments of different arithmetic intensities. These segments are then handled by separate kernels, with long segments aggregated and prioritized within the kernel. In comparison to AVX512 accelerated mm2-fast, our evaluations demonstrate that *mm2-gb* running on an AMD Instinct™ MI210 GPU achieves a 2.57-5.33x performance improvement on long reads (10kb-100kb) and up to a 1.87x performance gain on ultra long reads (100kb-300kb).

## 6 Related Works

A few prior works have attempted to speed up Minimap2. Zeni et al. [19] and Feng et al. [6] sped up the base-level alignment step, which is no longer the bottleneck as read lengths have increased. Some of the works that have accelerated chaining are either closed source [16], low in accuracy [8], or do not completely exploit the parallelism in chaining [10]. Guo et al. [8] introduced the concept of forward transforming the chaining algorithm and accelerated it on GPU and Field Programmable Gate Array (FPGA). However, their work does not guarantee output equivalence to mm2. Their decrease in mapping accuracy, as pointed out by Sadasivan et al. [16], is primarily due to Guo et al.’s static predecessor range selection, which differs from the dynamic selection in mm2, and because the rules for updating chaining scores were not modified in line with the forward transformation. Another study, mm2-fast [10], accelerates all three steps in mm2 using Single Instruction Multiple Data (SIMD) CPUs. While mm2-fast does parallelize the generation of chain scores, Sadasivan et al. [16] identified certain sections of chaining that are not parallelized. Sadasivan et al. [16] found that 34.08% of the total time spent generating chain scores and finding the maximum scores at the start and end of chains by mm2-fast is spent in sequential code and is not parallelized. mm2-ax [16] attempts to improve upon mm2-fast by forward transforming the chaining step while maintaining high accuracy. mm2-ax’s findings have motivated some of the strategies we have investigated in this work. Despite mm2-ax doing a good job at exploiting more parallelism in chaining, we identify several areas for improvement. mm2-ax’s workload scheduling is not designed to deliver consistent performance for longer read lengths, is fine-tuned for and requires NVIDIA A100 GPUs, is closed-source, and tied to an older version of mm2-v2.17. We compare with respect to mm2-fast because it is based on the latest mm2-v2.24, which is considered accurate for diagnostics.

## 7 Discussion

### 7.1 Latency Bounded Long Kernel

We profile the most time-consuming kernel of *mm2-gb*, the long score generation kernel on an ultra-long read ONT dataset (reads between 90kb-100kb extracted from 60x HG002 dataset) with Omniperf v1.0.10 [14]. As illustrated in Table 1, the long score generation kernel does not fully saturate L1, L2, or HBM memory bandwidth. However, as shown in Tab. 2, both the issue pipeline and vector ALU are under-utilized, with more than 70% of CU cycles being consumed by dependency stalls while awaiting data from the memory system. With a high L2 hit rate of 98%, a low L1 hit rate of 68%, and an average L1 latency of 216 cycles, we infer that the long score generation is constrained by L2 data fetch latency resulting from L1 capacity misses.

**Table 1.**
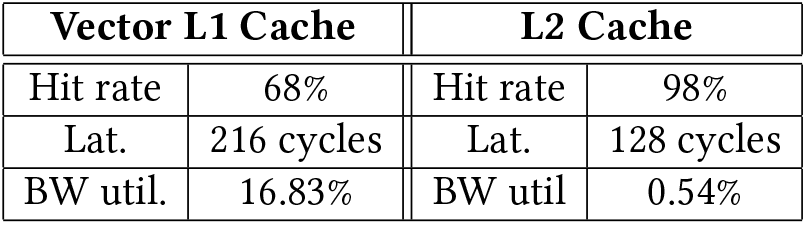
Cache statistics for long score generation kernels, as reported by omniperf.

**Table 2.** CU statistics for long score generation kernels, as reported by Omniperf.

| Compute Unit Utils |  | CU Cycle Breakdown |  |
| --- | --- | --- | --- |
| VALU Util | 32.57% | Dependency Wait | 73.50% |
| IPC-Issue Util | 19.91% | Issue Wait | 8.18% |
|  |  | Active Cycles | 18.93% |

### 7.2 Portability

The versatility of *mm2-gb* across different architectures and configurations is best highlighted by the following three aspects:

1. Cross-platform compatibility: *mm2-gb* is built on the Heterogeneous-compute Interface for Portability (HIP) developed by AMD. It runs on both AMD and NVIDIA GPUs.
2. Customization: We provide a user interface for fine-tuning kernel parameters via a JSON file. This allows achieving optimal performance on different hardware configurations.
3. Open Source: *mm2-gb* is open-source and hence can be easily modified to meet specific user requirements.

## 8 Future Work

### 8.1 Long Kernel: Shared Memory Prefetch

Our long score generation kernel is memory latency bound. However, we observe marginal benefits from L1 prefetching as opposed to mm2-ax. mm2-ax opted for L1 prefetching over shared memory prefetching due to limited temporal locality unified score generation kernel, whereas *mm2-gb*’s long kernel operates on aggregated long segments with extended predecessor ranges. These long-range segments exhibit greater data reuse between iterations, and the working set size of a block (estimated to be the size of anchors in the range) is also larger. For instance, in a segment with an average range of 3k anchors, the working set size of the block processing this segment is estimated to be 7*B*/*anchor* × 3*K*/*anchor* = 277*KB*. An AMD Instinct™ MI210 device can accommodate two long score generation blocks on one CU, sharing a 16KB L1 cache and a 64KB shared memory. The working set fits within the shared memory but exceeds the capacity of the L1 cache. Therefore, we anticipate that the long score generation kernel will benefit from shared memory prefetching, and we plan to implement it in our future updates.

### 8.2 Exploring the APU Architecture

The APU architecture in the AMD Instinct™ MI300A has some attractive features for accelerating minimap2 even further. With a larger HBM capacity of 128 GB and a unified memory addressing space, we can eliminate an additional copy of arrays and memory copies that would otherwise be required in traditional discrete memory space architectures such as MI210 GPUs. With the additional HBM capacity, we may be able to fit more batches of work and aggregate more long segments to achieve higher CU occupancy. Evaluation of mm2-gb on the APU architecture remains as future work.

### 8.3 Scalability

In the future, we want to explore scaling mm2-gb to support multiple CPU threads and using HIP streams to launch work on the GPU device concurrently. On a system with multiple GPUs, we intend to measure the throughput achieved when running multiple mm2-gb processes, each using a GPU device and processing a different set of reads.

## 9 Availability

### 9.1 Software

*mm2-gb* is open-sourced and available at https://github.com/Minimap2onGPU/mm2-gb.

### 9.2 Datasets

Datasets are publicly available with CC0 license from Human Pangenome Reference Consortium: https://github.com/human-pangenomics/HG002_Data_Freeze_v1.0 [5].

## Acknowledgments

We thank Ossian O’Reilly for valuable guidance on project direction and for being available for innumerable discussions, especially in the initial stages. We thank the capstone design program of Shanghai Jiao Tong University Joint Institute where this project was initiated, and Chen Huang, Xiang-dong Wei, and Haoyang Zhang who contributed to those initial discussions and deep dives into the alignment step of Minimap2. We are grateful to Ozzie Moreno and Anand Raghavendra of the AMD Accelerator Cloud (AAC) team for actively supporting us with computing infrastructure. This project was supported in part by AMD Data Center GPU (DCGPU), AMD University Program (AUP), the Kahn Foundation and NSF award 2403119.

## Footnotes

1 Intel is a trademark of Intel Corporation or its subsidiaries

2 Xeon is a trademark of Intel Corporation or its subsidiaries

